# LINC01184 is highly expressed and functions as a tumor-promotive factor in hepatocellular carcinoma

**DOI:** 10.1101/2022.10.09.511516

**Authors:** Qiangnu Zhang, Lesen Yan, Jiaojuan Chen, Liping Liu

## Abstract

The expression and function of LINC01184 in hepatocellular carcinoma are unknown. We analyzed the changes in LINC01184 expression in hepatocellular carcinoma using public data, and also analyzed the relationship between LINC01184 and patient prognosis. In vitro, we investigated the effect of LINC01184 on HCC cell growth using cell activity analysis, clone formation analysis, and regulatory death analysis. We also explored the effect of LINC01184 on HCC cell migration using transwell analysis and scratch assay. We found that LINC01184 was significantly highly expressed in hepatocellular carcinoma tissues. Patients with high LINC01184 levels had poorer overall survival and disease-free survival. knockdown of LINC01184 in HUH7 and PLC/PRF/5 cells inhibited cell proliferation and induced regulatory death. Down-regulation of LINC01184 also leads to a reduction in the migratory capacity of cells. In conclusion, LINC01184 may become a biomarker or a new therapeutic target for HCC.

## Introduction

Hepatocellular carcinoma (HCC) accounts for over 90% of primary liver cancer diagnoses and remains a poorly managed malignancy with high mortality. Approximately 0.9 milion new cases of HCC are diagnosed annually and 0.8 million deaths due to liver cancer worldwide, which were the sixth highest number of cases of cancer and the third highest number of deaths from cancer worldwide, respectively[1]. Despite great advances in the therapeutic strategies (e.g. immune-based intervention or targeted therapy) for HCC in the last decade, the long-term survival rate of HCC patients remains dismal[2]. Hence, the mechanisms that drive aggressive progression of HCC remain need to be reveal.

Long noncoding RNAs (lncRNAs) are a type of RNA molecules l longer than 200 nucleotides (nt) and with no or weak protein-coding capacity[3]. Increasing evidence has indicated that lncRNAs play roles in tumor including regulating prolifetration, apoptosis and stemness[4]. Therefore lncRNAs may become a new perspective to explain tumor development and have potential for clinical application in tumor diagnosis and treatment[5].

LINC01184 was found that obviously upregulated in colorectal cancer (CRC) tissues and cells and knockdown of LINC01184 significantly suppressed CRC cell proliferation and invasion and promoted apoptosis[6]. However, the expression characteristics and role of LINC01184 in HCC are not clear. We analyzed the expression of LINC01184 in HCC in this study and explored its function in a cellular model.

## Materials and Methods

### Public data collection

The RNA sequencing (RNA-seq) data and clinical data of The Cancer Genome Atlas project (TCGA-LIHC) were collected from UCSC Xena public datahub (http://xena.ucsc.edu/public/). The gene tissue specificity index of lncRNAs were obtained from Human Ageing Genomic Resources (https://genomics.senescence.info/gene_expression/tau.html). The Pan-cancer data of LINC01184 were obtained from GEPIA database (http://gepia.cancer-pku.cn/).

### Cell culture

HUH7, PLC/PRF/5 were obtained from the Cell Bank of the Chinese Academy of Sciences. All cell lines were cultured in Dulbecco’s modified Eagle’s medium (DMEM, Gibco, CA, USA) with 10% fetal bovine and 1% penicillin/streptomycin in 5% CO_2_ at 37°C.

### Cell viability assay

Transfected or treated cells were planted into a 96-wells plate at a density of 3000 cells/100 μL. CellTiter-Lumi™ Steady Plus Luminescent Cell Viability Assay Kit was used to detect the cells viability changes at multiple time points. 100 μL detection reagent was added for each well. The luminescence intensity was detected by a luminometer (SPARK 10M) after 10 min incubation.

### Colony formation

The indicated number of cells were planted in a 6-well plate and cultured in an incubator for 3-10 days. Colonies c were stained with 0.1% crystal violet dye and subsequently counted.

### Healing assay assay

For wound healing assay, 2-Well ibidi Culture-Inserts were used and 70 μL cells (30 0000cells/mL) were planted for each well. The gap was made after 24 h and images were obtained at indicated time point.

### Analysis of apoptosis

The apoptosis of treated cells was continuously monitored by the RealTime Glo Annexin V Apoptosis Assay kit. In brief, 5000 cells were planted into a 96-well plate in 100 μL DMEM. 0.1 μL Annexin V NanoBiT™ Substrate, 0.1 μL Annexin V-SmBiT, and 0.1 μL Annexin V-LgBiT were added to each well. Luminescence was read by a luminometer (SPARK 10M).

### Statistical analysis

R 4.1.0 software (https://www.R-project.org/) and GraphPad Prism 7.0 (GraphPad Software, CA, USA) was used for statistical analysis. Data are presented as mean ± standard deviation. Differences between two different groups were evaluated using a t-test. P < 0.05 was defined as statistical significance.

## Results

### High expression of LINC01184 is associated with poor prognosis in patients with HCC

In the TCGA-LIHC data (Figure 1A) and the TCGA-LIHC+GTEx data (Figure 1B), LINC01184 was significantly highly expressed in HCC tissues (P<0.05), compared with normal tissues. However LINC01184 levels was not associated with the progression of TNM staging (Figure 1C). We divided patients from TCGA-LIHC into a high LINC01184 group and a low LINC01184 group using the median value of LINC01184 as the cut-off value. K-M survival analysis found that patients in the high LINC01184 group had poorer overall survival (Figure 1D) and disease-free survival (Figure 1E).

**Figure 1.**
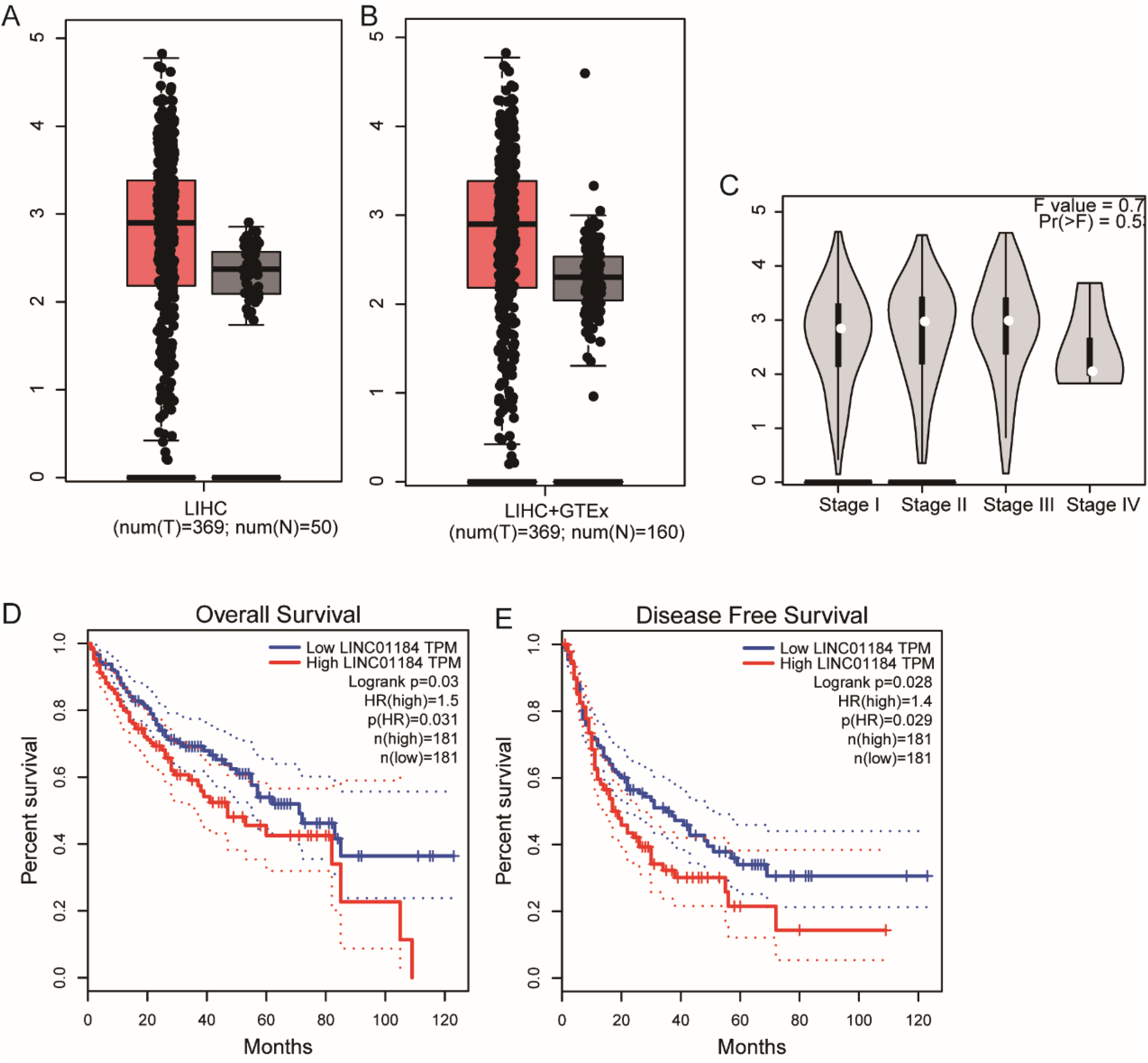
LINC01184 highly expressed in HCC and indicated poor prognosis. (A-B) LINC01184 levels in HCC and normal tissues from TCGA-LIHC and GTEx datasets; (C) LINC01184 levels in HCC patients with different TNM stages;(D-E) High LINC01184 indicated poor overall survival and disease-free survival.

### Knockdown of LINC01184 inhibites progression of HCC cells in vitro

Knockdown of LINC01184 significantly reduced the cellular activity of HUH7 and PLC/PRF/5 cells (Figure 2A). The colony formation assay suggested that knockdown of LINC01184 could inhibit the proliferation of HCC cells (Figure2B). Real-time apoptosis analysis revealed that reducing of LINC01184 could induce apoptosis in HCC cells (Figure 2C). Transwell analysis suggested that downregulated of LINC01184 could inhibit the migration of HCC cells (Figure 2D). Similar results were obtained by scratch assay (Figure2E).

**Figure 2.**
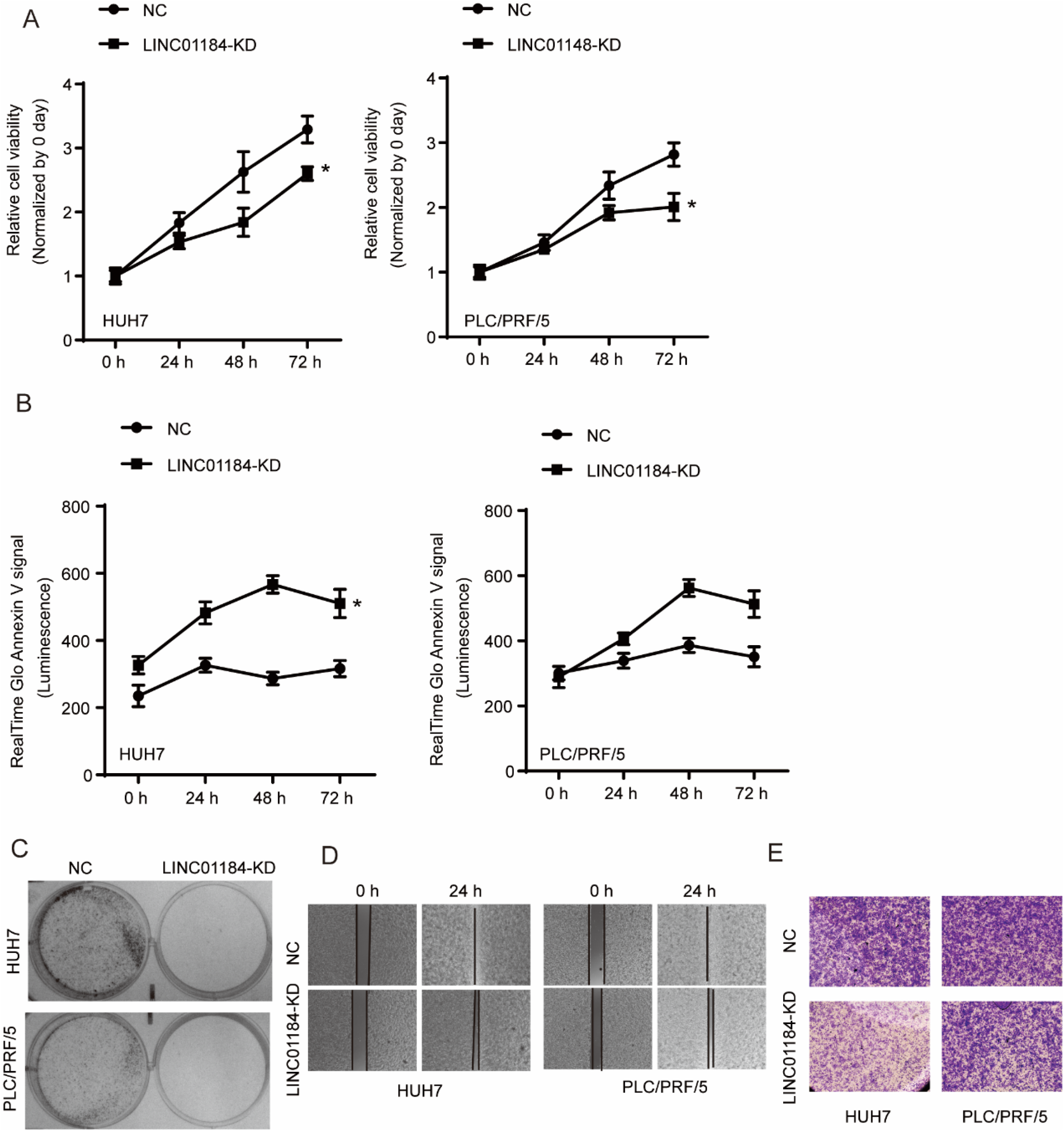
Knockdown of LINC01184 inhibits progression of HCC in HUH7 and PLC/PRF/5 cells. (A) CCK-8 assay was used to estimate the cell viability changes after LINC01184 knockdown; (B) Clone formation assay for HUIH7 and PLC/PRF/5 cells with LINC01184 knockdown; (C) Real-time apoptosis assay for HUIH7 and PLC/PRF/5 cells with LINC01184 knockdown; (D-E) Transwell assay and wound healing assay for HUIH7 and PLC/PRF/5 cells with LINC01184 knockdown *P<0.05.

## Discussion

In the present study, we analyzed the expression characteristics of LINC01184 in hepatocellular carcinoma, and we found that LINC01184 was highly expressed in hepatocellular carcinoma, and patients with high expression of LINC0118 had poor survival. These data suggested that LINC01184 plays a role in promoting tumor growth in hepatocellular carcinoma. We subsequently confirmed this hypothesis using an *in vitro* cell model, where we found that knockdown of LINC01184 resulted in slower growth of HUH7 and PLC/PRF/5 cells, along with additional regulatory death. Knockdown of LINC01184 was also able to inhibit the migration of both HCC cells.

The role, function, and mechanism of LINC01184 has been already reported by researchers. LINC01184 was found that up-regulated in CRC and indicated poor outcome. Mechanically, LINC01184 regulated miR-331/ HER2 pathway and hence controlled AKT and ERK signaling[6]. As a kind of non-coding RNA that has received much attention in recent years, the mechanism of its function, in addition to ceRNA-related mechanisms, has also been continuously revealed. Among the many mechanisms that have been discovered, it is mainly dependent on the RNA molecule itself, such as acting as a molecular scaffold[7], a molecular signal[8], a capture molecule or a mediator to participate in the regulation of gene expression and function[9, 10]. We have not yet discovered the mechanism by which LINC01184 plays a role in hepatocellular carcinoma, and it is just an interesting question that needs to be addressed in the future.

In conclusion, we identified LINC01184 as a potential cancer-promoting gene in hepatocellular carcinoma. It might be used as a diagnostic marker or a therapeutic target for HCC.

